# Active behaviour during early development shapes glucocorticoid reactivity

**DOI:** 10.1101/582163

**Authors:** Luis A. Castillo-Ramírez, Soojin Ryu, Rodrigo J. De Marco

**Author notes:** **Author for correspondence:** E-mail: R.J.D.M. < >.

## Abstract

Glucocorticoids are the final effectors of the stress axis, with numerous targets in the central nervous system and the periphery. They are essential for adaptation, yet currently it is unclear how early life events program the glucocorticoid response to stress. Here we provide evidence that involuntary swimming at early developmental stages can reconfigure the cortisol response to homotypic and heterotypic stress in larval zebrafish (*Danio rerio*), also reducing startle reactivity and increasing spontaneous activity as well as energy efficiency during active behaviour. Collectively, these data identify a role of the genetically malleable zebrafish for linking early life stress with glucocorticoid function in later life.

## Introduction

The increased secretion of glucocorticoids like cortisol after the onset of stress (a.k.a.) glucocorticoid reactivity (GC_R_) plays a pivotal role in the response to challenge. It is critical for adaptation and central to an organism’s resilience^1^. GC_R_ is a tightly regulated phenomenon, a response of the hypothalamic-pituitary-adrenal (HPA) axis to exogenous or endogenous stressors. Altered functionality of the HPA axis and of GC_R_ have been associated with detrimental and beneficial consequences for health. They have been linked to stress-evoked disorders including mental disorders as well as increased resilience^2,3,4,5,6,7,8^. Glucocorticoid secretion has been investigated extensively under steady-state and stress conditions^9^, and there is ample evidence that HPA axis functionality is susceptible to disturbance by early life stress. Early adversity can, for example, alter glucocorticoid regulation and coping capacities later in life^10,11^. However, it is still unclear how active responses to early life stress can reconfigure HPA axis function, pending a detailed functional evaluation of developmental programming of GC_R_. Larval zebrafish are excellent to address this knowledge gap due to their external development, their hypothalamic-pituitary-interrenal (HPI) axis, homologous to the mammalian HPA axis^12^, their translucent body, ideal for non-invasive brain imaging and optogenetics^13,14,15^, their small size, highly suitable for high-throughput screens with full environmental control, and the availability of tools and methods for identifying genetic and epigenetic modulators, including proteomic technology. Therefore, as a first step, we set out to determine the effect of early life stress on GC_R_ and coping capacity in larval zebrafish. Taken together our results introduce a high-throughput forced swim test for developing zebrafish and demonstrate that mild early life stress can at least transiently reconfigure GC_R_ and elicit modulatory adjustments in spontaneous activity and startle reactivity.

## Results

### High-throughput induction of forced swimming and cortisol increase

Firstly, we exposed groups of larvae to water vortex flows of varying strength, expressed in revolutions per minute (rpm) (for details, see the ‘Methods’ section). To compare the strength of these flows, we video-recorded and examined the paths (x-y coordinates) of anesthetized larvae (i.e., unable to swim) exposed to vortex flows of increasing rpm (Fig. 1a). The results of these observations confirmed that, as rpm increased, anesthetized larvae followed the vortex currents, thereby moving at higher speeds and larger distances from the source of the vortex. These measurements were used to determine vortex flows of low, medium and high strength. We then assessed the relationship between the strength of the flows and the behaviour of freely swimming larvae. Larval zebrafish have been shown to display positive rheotaxis^16,17^, i.e., spontaneous swimming against an oncoming current, which allows them to hold their position instead of being swept downstream by the current. When exposed to vortex flows, freely swimming larvae held their position away from the vortex’s source, thereby avoiding the strongest currents (Fig. 1b, Kruskal-Wallis test, H=27.6, *p* < 0.0001, followed by Dunn’s multiple comparison tests). They also faced the oncoming current (Fig. 1c, top, Chi-square test, X^2^(1, N=270)=55.4, *p* < 0.0001) and adjusted their swim bouts and turns (Fig. 1c, bottom, one-way ANOVA, F(2,29)=6.7, *p* = 0.004, followed by *post hoc* comparisons) to compensate for vortex strength. As indicated by their GC_R_, these behaviours were taxing for the larvae. Their whole-body cortisol increased together with the strength of the vortex (Fig. 1d, one-way ANOVA, F(2,17)=35.4, *p* < 0.0001, followed by *post hoc* comparisons).

**Figure 1.**
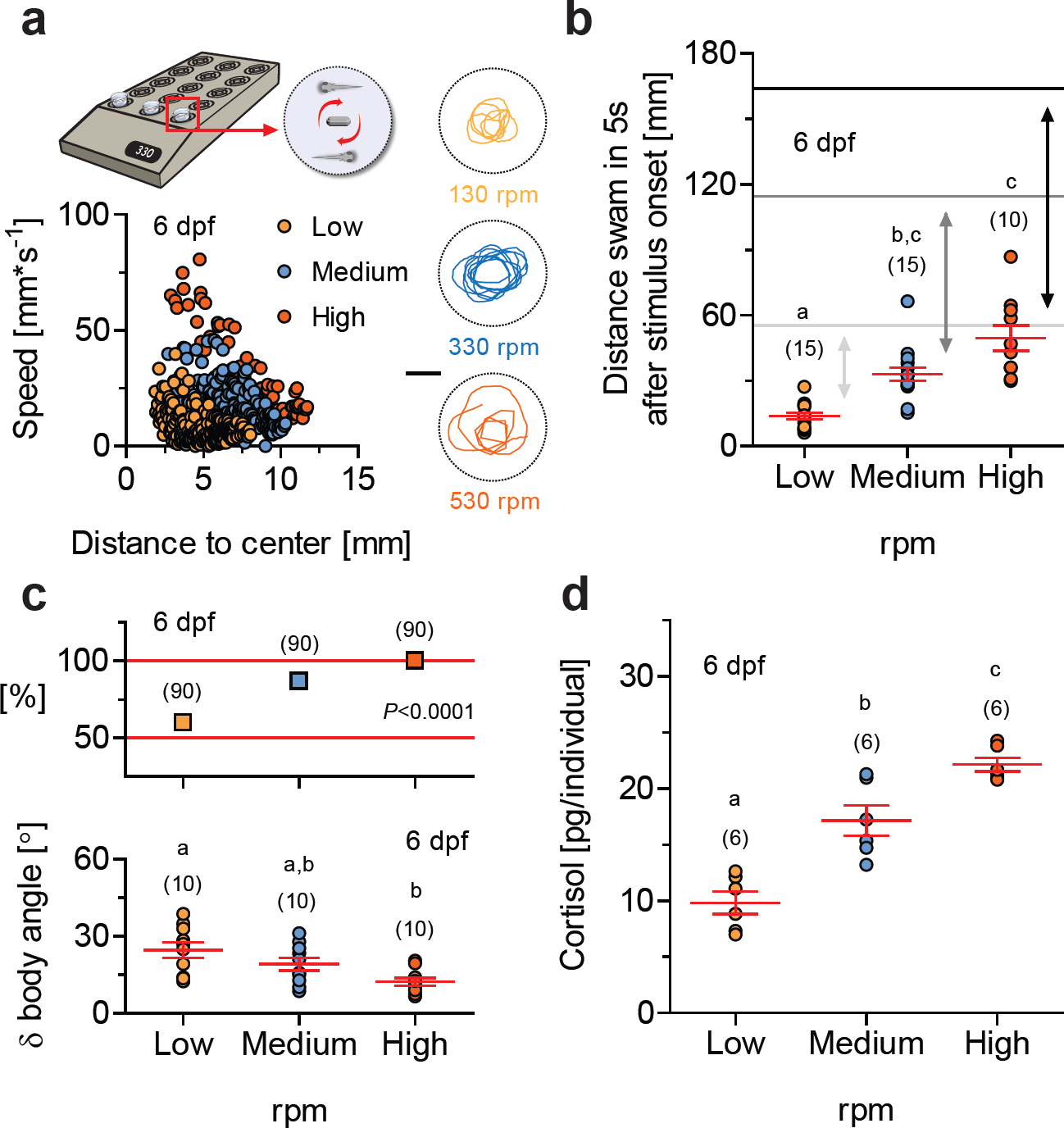
High-throughput induction of forced swimming and cortisol increase. (**a**) Top, schematic showing how groups of thirty zebrafish larvae in 35 mm diameter petri dishes can be exposed to water vortex flows in a high-throughput manner while their behaviour is being video-recorded (see the ‘Methods’ section). Right, representative x-y coordinates (recorded every 93.3 ms over a 2.4 s period) of single anesthetized 6 dpf larvae exposed to vortex flows of increasing strength, expressed in revolutions per minute (rpm). Scale bar, 10 mm. Bottom, swim velocity and distance to the center of the dish of a larva exposed to vortex flows of low (orange), medium (blue) and high (vermilion) strength levels (data from the top figures). (**b**) Distance swam in 5 s by freely behaving larvae after the onset of vortex flows as a function of vortex strength (as in **a**). Grey and black lines indicate the average distance covered by anesthetized larvae under similar conditions; double headed arrows highlight the differences between anesthetized and freely behaving larvae due to rheotaxis: the higher the vortex strength the lower the distance covered by individuals engaged in rheotaxis. Letters indicate results of Dunn’s multiple comparison tests (*p* < 0.01) after a Kruskal-Wallis test. (**c**) Top, Proportion of larvae engaged in rheotaxis (measured 120 s after the onset of vortex flows) as a function of vortex strength (as in **a**); *P*<0.0001 after a Chi-square test. Bottom, Average change in orientation after a swim bout (δ body angle, in degrees, recorded every 933 ms over a 5 s period 120 s after the onset of vortex flows) of freely swimming larvae as a function of vortex strength (as in **a**). (**d**) Whole-body cortisol in 6 dpf larvae as a function of vortex strength (as in **a**). (**c,d**) Letters indicate results of Bonferroni’s tests (*p* < 0.01) after one-way ANOVAs. (**b,c,d**) Sample size in parentheses.

### Cortisol change in response to water vortex flows as a function of development

Secondly, we assessed GC_R_ to vortex flows as a function of development, expressed in days post fertilization (dpf). For these tests we selected vortex flows of medium strength (i.e., 330 rpm), avoiding the high strength causing maximum levels of vortex-dependent cortisol increase and occasional disruptions of positive rheotaxis (not shown). The HPI axis of zebrafish matures early. Basal whole-body cortisol and expression levels of genes involved in corticosteroid synthesis and signaling increase drastically around the time of hatching^18,19^. We observed that whole-body cortisol increased gradually between 2 and 8 dpf (Fig. 2a, top, Kruskal-Wallis test, H=64.7, *p* < 0.0001, followed by Dunn’s multiple comparison tests), and that the magnitude of the vortex-dependent elevation of cortisol peaked at 6 dpf (Fig. 2a, bottom, Kruskal-Wallis test, H=36.5, *p* < 0.0001, followed by Dunn’s multiple comparison tests), with circulating levels of cortisol measured ten minutes after a three minute exposure to vortex flows (for details, see the ‘Methods’ section). Mechanistically, this points to fundamental alterations in the HPI axis occurring at 4-6 dpf.

**Figure 2.**
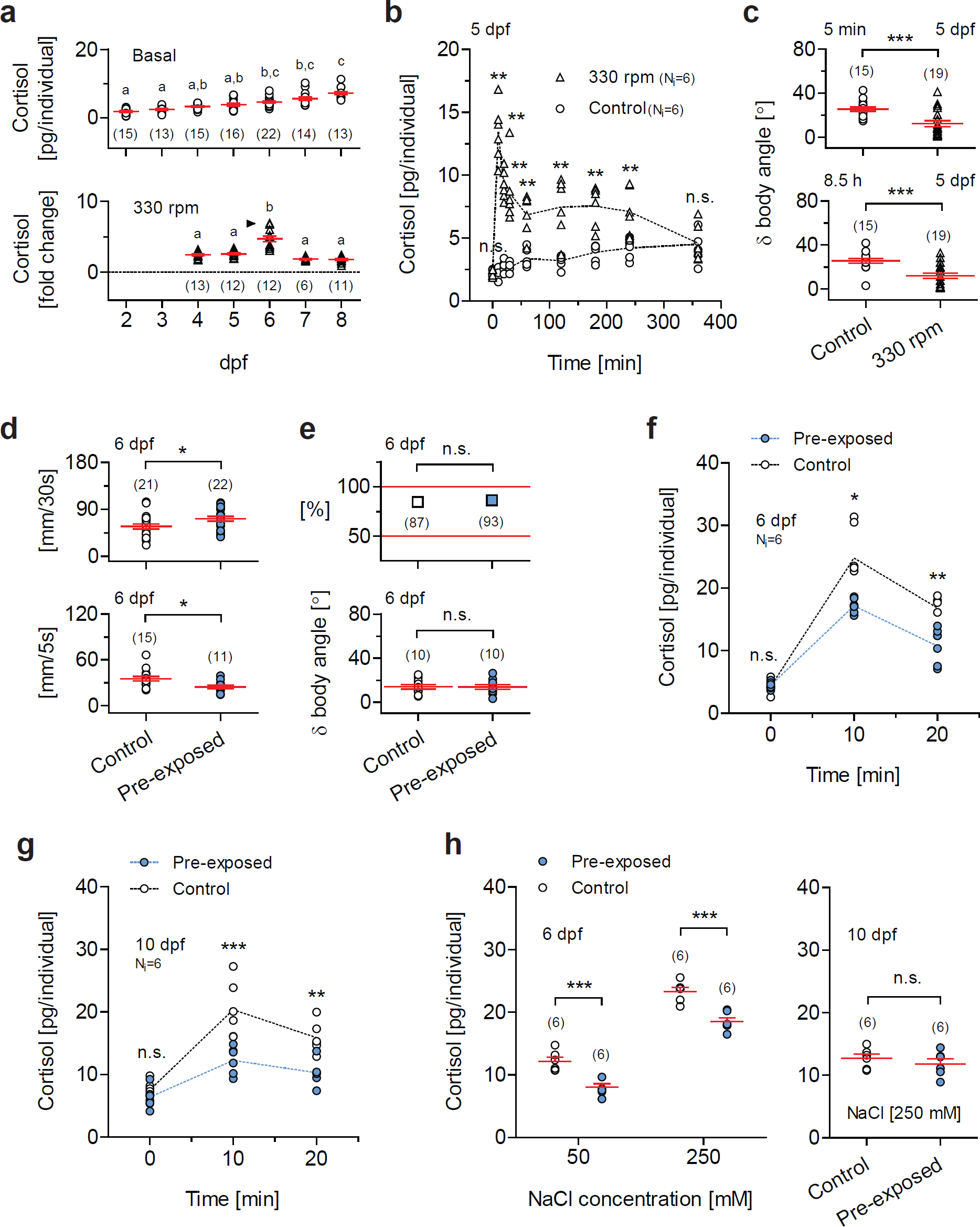
Prolonged forced swimming during early development increases spontaneous activity and reduces startle and glucocorticoid reactivity. (**a**) Top, Basal cortisol as a function of time, expressed in days post fertilization (dpf). Bottom, Cortisol change in response to vortex flows of medium strength (330 rpm) as a function of development, in dpf. Letters indicate results of Dunn’s multiple comparison tests (*p* < 0.01) after Kruskal-Wallis tests. (**b**) Cortisol time course in 5 dpf larvae exposed to vortex flows (330 rpm) for 9 hours (shown up to 6 hours) and controls (unexposed larvae). Cortisol in exposed larvae peaks shortly after the onset of the vortex and remains high 4 hours later; exposed and control larvae show similar values 6 hours after the onset of the vortex. (**c**) δ body angle (as in **Fig. 1c**), indicative of rheotaxis, in exposed and control 5 dpf larvae, measured 5 minutes (top) and 8.5 hours (bottom) after the onset of the vortex (330 rpm). \*\**P*<0.01, \*\*\**P*<0.001 after two-tailed *t*-tests. (**d**) Top, spontaneous activity (in mm swam in 30s) in pre-exposed and control 6 dpf larvae. Bottom, locomotor reaction to the onset of the vortex (330 rpm) (mm swam in 5s, measured 5 s after the onset) in pre-exposed and control 6 dpf larvae. Pre-exposed larvae, blue. Control larvae, white. *P*=0.03 (top) and *P*=0.01 (bottom) after two-tailed *t*-tests. (**e**) Proportion of individuals engaged in rheotaxis (top) and δ body angle (bottom) (as in **Fig. 1c**) in pre-exposed (blue) and control (white) 6 dpf larvae. Top, *P*=0.24 after a Chi-square test. Bottom, *P*=0.97 after a two-tailed *t*-test. (**f,g**) Cortisol in pre-exposed (blue) and control (white) 6 dpf (**f**) and 10 dpf (**g**) larvae, before, 10 and 20 minutes after the onset of the vortex. (**h**) Cortisol in pre-exposed (blue) and control (white) 6 dpf (left) and 10 dpf (right) larvae in response to a 10 min incubation in hyperosmotic medium. Right, *P*=0.42 after a two-tailed *t*-test. (**b,f**) Asterisks (\**P*<0.05, \*\**P*<0.01) indicate results of two-group comparisons using Mann-Whitney or two-tailed *t*-tests after Kruskal-Wallis tests. (**g,h**) Asterisks (\*\**P*<0.01, \*\*\**P*<0.001) indicate results of Bonferroni’s tests after two-way ANOVAs. (**b,f,g**) Sample size per group, 6. (**a,c,d,e,h**) Sample size in parentheses.

### Prolonged forced swimming and HPI activation at 5 dpf

Thirdly, building on the above findings, we exposed 5 dpf larvae to vortex flows of medium strength for 9 hours. Exposed larvae showed increased levels of whole-body cortisol which peaked shortly after the onset of the vortex and remained high four hours later compared with controls, i.e., unexposed larvae that were equally handled, but the vortex flows were not present. Both exposed and control larvae showed similar levels of whole-body cortisol six hours after the onset of the vortex (Fig. 2b, Kruskal-Wallis tests, exposed larvae: H=36.9, *p* < 0.0001, control larvae: H=36.3, *p* < 0.0001, followed by two-group comparisons using Mann-Whitney tests). Also, exposed larvae remained engaged in positive rheotaxis from the beginning to the end of the vortex, as indicated by their δ body angle (average change in orientation after a swim bout) measured 5 minutes (Fig. 2c, top, two-tailed *t*-test, t(32)=3.8, *P* = 0.0007) and 8.5 hours (Fig. 2c, bottom, two-tailed *t*-test, t(32)=4.1, *P* = 0.0003) after the onset of vortex flows (see also the ‘Methods’ section).

### Increased baseline swimming and reduced startle reactivity in pre-exposed larvae

Next, using video-recordings and off-line measurements (for details, see the ‘Methods’ section), we assessed the behaviour of 6 dpf larvae that had or had not been exposed to vortex flows at 5 dpf, i.e., pre-exposed and control larvae, respectively. Compared with controls, pre-exposed larvae showed higher levels of baseline swimming (Fig. 2d, top, two-tailed *t*-test, t(41)=2.2, *P* = 0.03) and reduced startle reactivity upon re-exposure to vortex flows of medium strength, as specified by the distance they swam directly after the onset of the water current (Fig. 2d, bottom, two-tailed *t*-test, t(24)=2.7, *P* = 0.01). Importantly, pre-exposed larvae engaged in positive rheotaxis as efficiently as controls, as indicated by the proportion of larvae facing the oncoming current (Fig. 2e, top, Chi-square test, X^2^(1, N=180)=1.4, *p* = 0.24) and δ body angle (Fig. 2e, bottom, two-tailed *t*-test, t(18)=0.08, *P* = 0.94).

### Reduced glucocorticoid reactivity to vortex flows in pre-exposed larvae

At 6 dpf, pre-exposed larvae, which had prior experience with the vortex flows at 5 dpf, displayed the above behavioural adjustments as well as reduced GC_R_. Relative to controls, pre-exposed larvae showed similar levels of basal cortisol and reduced levels of vortex-dependent cortisol increase upon re-exposure to vortex flows of medium strength (Fig. 2f, Kruskal-Wallis tests, pre-exposed larvae: H=15.2, *p* = 0.0005, control larvae: H=13.7, *p* = 0.001, followed by two-group comparisons using two-tailed *t*- or Mann-Whitney tests, basal cortisol: t(10)=0.48, *P* = 0.65, 10 minutes after exposure: U=2, *P* = 0.01, 20 minutes after exposure: t(10)=4.0, *P* = 0.003). We found the same pattern of results at 10 dpf (Fig. 2g, two-way ANOVA, group: F(1,30)=32.8, *p* < 0.0001, time: F(2,30)=40.5, *p* < 0.0001, group × time: F(2,30)=5.6, *p* = 0.009, followed by *post hoc* comparisons).

### Short-term reduced glucocorticoid reactivity to heterotypic stress

To complement these assessments we examined the relationship between GC_R_ in pre-exposed larvae and heterotypic stress. For this we exposed 6 dpf pre-exposed and control larvae to hyperosmotic medium (NaCl), a known stress protocol (for details, see the ‘Methods’ section). The results showed that, relative to controls, pre-exposed larvae showed reduced GC_R_ to moderate and high levels of salt stress (Fig. 2h, left, two-way ANOVA, group: F(1,20)=51.0, *p* < 0.0001, NaCl concentration: F(1,20)=299.0, *p* < 0.0001, group × NaCl concentration: F(1,20)=0.3, *p* = 0.59, followed by *post hoc* comparisons). By contrast, at 10 dpf, both groups showed similar cortisol responses to salt stress (Fig. 2h, right, two-tailed *t*-test, t(10)=0.9, *P* = 0.42).

## Discussion

Hormones react to the environment and cause changes in physiology as a function of maturation. Thus the question arises as to how dynamical patterns of hormone secretion are achieved and what effects they exert on well-being. How does the environment activate and guide the development of resilience mechanisms? Current paradigms stipulate that glucocorticoids are fundamental to the mitigation of allostatic load^9,20^. However, the impact of early life stress on developmental programming of GC_R_ has not been explored in full, in part due to a lack of suitable models. We now show in zebrafish that the increase in cortisol elicited by a brief period of involuntary swimming peaks at 6 days post fertilization (dpf), pointing to fundamental changes in GC_R_ at early larval stages. Importantly, we found that, if prolonged for hours, forced swimming at 5 dpf caused a transient form of hypercortisolaemia and later led to reduced GC_R_. If subsequently exposed to a brief period of involuntary swimming (i.e., homotypic stress), pre-exposed larvae showed a decreased cortisol response that persisted for at least five more days. Moreover, twenty-four hours after prolonged forced swimming at 5 dpf, the reduced GC_R_ appeared invariant to stressor identity, as indicated by a decreased cortisol response to heterotypic stress, i.e., osmotic shock. These data suggested that the sustained reduction in GC_R_ did not reflect a process of habituation to sensory input^21^. It seems likely that, in pre-exposed larvae, reduced GC_R_ was underpinned by changes in state variables of the HPI axis.

Early stress as well as chronic stress in later life can decrease hypothalamic activity and expression of corticotropin-releasing-hormone (CRH) and arginine-vasopressin (AVP). These changes are mediated at least in part by glucocorticoids^22,23,24,25^. Previous studies in fish showed that prolonged stimulation of the HPI axis can attenuate the stress response^26^. This effect can result from transcriptional regulation of CRH and adrenocorticotropic hormone (ACTH)^27,28,29^. Additionally, pituitary corticotrophs and cortisol-producing cells in the interrenal gland may be desensitized to CRH or ACTH, respectively^30,31^. Stressor exposure on early stages of rainbow trout can lead to HPI axis hypoactivity later in life^32^. In zebrafish, incubation in cortisol during the first 48 hours post fertilization caused altered locomotor reactions to photic stimuli^33^. Also, incubating zebrafish embryos in cortisol during the first five days post fertilization caused increased whole-body cortisol, glucocorticoid signalling and expression of immune-related genes; these changes can be long-lasting and result in dysfunctional capacities for regeneration along with increased expression of inflammatory genes^34^. A similar treatment using dexamethasone also induced long-lasting behavioural and metabolic changes still detectable in adulthood^35^. Further experiments are necessary to determine whether reduced GC_R_ in pre-exposed larvae occurs via receptor downregulation, decreased synthesis and/or depletion of hormones, and/or increased sensitivity to glucocorticoid feedbacks^36,37,38^.

An organism is said to be engaged in active behaviour when it is the source of the output energy required for a given action^39^. Glucocorticoids are known to mobilize energy^1^, necessary to face the demands of forced swimming. In response to the vortex, larvae engaged in rheotaxis had to adjust their swim bouts and turns continuously to compensate for the oncoming current. These actions were energy demanding for the larvae, as revealed by their GC_R_. The notion that upholding positive rheotaxis for hours involved mobilizing energy was supported by the long-lasting hypercortisolic state observed during prolonged forced swimming. Twenty-four hours after prolonged exposure to the vortex, we observed differences between pre-exposed and control larvae. Firstly, pre-exposed larvae displayed increased levels of baseline swimming. A previous study in zebrafish showed that involuntary swimming at larval stages can subsequently increase spontaneous activity^40^. Secondly, upon a brief re-exposure to the same vortex, pre-exposed larvae showed reduced startle reactivity to the onset of water motions. Thirdly, they engaged in positive rheotaxis as efficiently as controls. On the assumption that the cortisol response to the vortex reflects an energy requirement for positive rheotaxis, these observations indicated that pre-exposed larvae responded more efficiently to the energy demands of forced swimming.

In conclusion, we have shown in larval zebrafish that early life stress caused by prolonged forced swimming at least transiently reconfigures the increased secretion of cortisol after the onset of homotypic or heterotypic stress, as well as spontaneous activity and efficient energy use during active behaviour. It remains open how these changes relate to survival in a species facing greater mortality during early life; there is a lack of evidence linking early activity patterns of the HPI axis to survival and reproductive outcome. Collectively, our data provided direct evidence to support the contention that long-term changes in HPI axis function after early adversity may later lead to increased resilience in developing zebrafish. Increased resilience to stress may have advantages for larval zebrafish. Such ability may help larvae to better cope with antagonistic environments. An important question emerging relates to the study of stress reactivity during adulthood as a function of early life events in zebrafish. A previous study in mice reported that individuals that had endured early life stress coped better with forced swimming compared with those that had experienced a favourable early care regime^41^. In adult rats, the adverse experience of maternal separation during early life strengthened freezing during fear conditioning after chronic stress compared with non-maternally separated rats^42^. These studies support the view that early life stress can subsequently lead to increased resilience. Long-lasting changes in HPA axis function due to early experiences have been attributed to changes in the epigenome^43^, although the link between early life stress and the activation of resilience mechanisms has been difficult to pin down in models with intrauterine development. In zebrafish, all three elements of the HPI axis can be visualized and genetically manipulated at early developmental stages and measured with modern molecular tools^13^. Moreover, the larval brain is readily accessible and provides excellent access for assessing how systematic variations in physiological and behavioural schemes relate to differences in the activity of neuronal and humoral networks. Further studies are required to determine the applicability of our high-throughput procedure. In anticipation to these studies, we speculate that zebrafish larvae will prove fruitful to link early HPI axis activity to proteomic regulation, epigenetic programming and measures of stress resilience.

## Methods

### Zebrafish husbandry and handling

Zebrafish breeding and maintenance were performed under standard conditions^44^. Groups of thirty wild-type embryos (cross of AB and TL strains, AB/TL) were collected in the morning and raised on a 12:12 light/dark cycle at 28 °C in 35 mm Petri dishes with 5 ml of E2 medium. At 3 days post fertilization (dpf), the E2 medium was renewed and chorions and debris were removed from the dishes. Experiments were carried out with 5-6 dpf larvae, with the exception of the cortisol measurements in Fig. 2a, g and h(left). Larvae older than 6 dpf were transferred to plastic cages with 400 ml of egg water in groups of thirty and fed with paramecia daily. Tests were performed between 09:00 hours and 18:00 hours, with different experimental groups intermixed throughout the day. Zebrafish experimental procedures were performed according to the guidelines of the German animal welfare law and approved by the local government (Regierungspräsidium Karlsruhe; G-29/12).

### Water vortex flows

Water current can trigger rheotaxis in larval zebrafish and, if sufficiently strong, it can also act as a stressor, causing a sharp increase in whole-body cortisol via the activation of the HPI axis. We used water vortex flows in a high-throughput fashion to induce both rheotaxis and cortisol increase. For this we exposed groups of thirty (4-8 dpf) larvae in 35 mm Petri dishes with 5 ml of E2 medium to the vortex flows caused by the spinning movements of small magnetic stir bars (6 × 3mm, Fischerbrand, #11888882, Fisher scientific, Leicestershire, UK.) inside the dishes. The Petri dishes, each with a single stir bar, were positioned on magnetic stirrer plates (Variomag, Poly 15; Thermo Fisher Scientific, Leicestershire, UK) and kept at 28°C inside an incubator (RuMed 3101, Rubarth Apparate GmbH, Laatzen, Germany). Larvae were presented with either short (3 minutes) or long (9 hours of continuous stimulation) exposure periods to the vortex flows caused by the highly-controlled magnetic field inversions of the stirrer plate, of 130, 330 or 530 revolutions per minute (rpm). For the short exposure, we avoided exposure periods longer than 3 minutes to elude maximum levels of stressor-mediated cortisol increase (not shown). The long exposure at 5 dpf consisted of 9 hours to achieve the longest possible exposure period adjustable to the light/dark cycle. Once exposed, larvae were immobilized in ice water and used for cortisol measurement (see below). Control larvae were collected after equal handling, omitting exposure to vortex flows (i.e., stir bars inside the Petri dishes were absent). To rule out unspecific effects of the magnetic field inversions produced by a stirrer plate, we compared the level of basal whole-body cortisol across groups of 6 dpf larvae that either remained unexposed or had been exposed to magnetic field inversions alone (i.e., without stir bars inside the Petri dishes and thus in the absence of vortex flows), of 130, 330 and 530 rpm. The results of these tests showed that magnetic field inversions per se did not alter the level of whole-body cortisol (one-way ANOVA, F(3,23) = 0.05, *p* = 0.98).

### Re-exposure to vortex flows

For these tests we selected vortex flows of medium strength to avoid possible ceiling effects caused by maximum levels of vortex-dependent cortisol increase. Using the above protocol, larvae that had or had not been exposed to vortex flows for 9 hours at 5 dpf were re-exposed to vortex flows (330 rpm) for 3 minutes at either 6 or 10 dpf. They were subsequently used for cortisol detection or behaviour evaluation.

### Re-exposure to vortex flows at 10 dpf

A plastic cage (5 L) containing thirty pre-exposed or control 10 dpf larvae and three magnetic stir bars (25 × 6 mm, Fisherbrand, #10226853, Fisher scientific, Leicestershire, UK) distributed equidistantly along the bottom of the cage were placed on top of the magnetic stirrer plate (Variomag, Poly 15, Thermo Scientific, Leicestershire, UK). Larvae were then exposed to vortex flows (330 rpm) for 3 minutes. Larvae were then immobilized with ice water and used for cortisol extraction 10 minutes after the onset of the vortex flows.

### Hyperosmotic medium

Groups of thirty larvae (either 6 or 10 dpf) in 35 mm Petri dishes were incubated for 10 min in steady state E2 medium (controls) or E2 + 50 or 250 mM NaCl (Merck, #106404, Darmstadt, Germany) at 28°C under white light illumination. They were washed three times with E2 medium and kept for immediate cortisol detection. The wash and transfer period took 3 min (± 10 s) and was performed at room temperature.

### Whole-body cortisol

The procedures and the home-made ELISA used for cortisol measurements were as previously described^13,45^. Groups of thirty larvae were immobilized in ice water after being exposed to water vortex flows or NaCl. Unexposed larvae (control samples) were collected after equal handling, omitting stressor exposure. Samples were then frozen in an ethanol/dry-ice bath and stored at −20 °C for subsequent extraction. Each replicate consisted of a well with 30 larvae. Cortisol extraction and detection were carried between 10:30 and 11:30 hours.

### Independent sampling

The scheme of independent sampling has been described elsewhere^46^. Briefly, cortisol and behavioural measurements were made on different groups of equally treated larvae and therefore constitute fully independent samples. For the behavioural measurements, each replicate involved a single larva. Yet, these individual measurements were made on larvae that had also been kept in wells containing a total of thirty larvae per well. Thus, the number of single larvae matched the number of independent wells. In this manner, the density of larvae per well during vortex flow exposure remained a constant factor for both the cortisol and behavioural measurements. For each cortisol measurement, all thirty larvae in a well were used, whereas each behavioural measurement involved only one larva, the remaining twenty-nine larvae in the well were used elsewhere. Each replication was fully independent from the others thus avoiding pseudo-replication.

### Anesthetized larvae

To assess the speed and trajectories of anesthetized larvae exposed to vortex flows of increasing strength, 6 dpf larvae in 35 mm Petri dishes were first incubated in 5 mL of steady state E2 medium + 100 µL of Tricaine (Sigma-Aldrich #E10521, Schnelldorf, Germany); they were considered to be anesthetized when they failed to respond to tactile stimulation. They were then transferred to a new Petri dish with fresh E2 medium (5 mL) for testing.

### Behaviour evaluation

Video recordings were conducted under conditions identical to those of the cortisol measurements. Groups of thirty larvae (either 5 or 6 dpf, depending on the experiment) were imaged at 12.5 frames s^−1^ with a camera (HDR-CX240 HD Flash, Sony, Berlin, Germany) positioned above a 35 mm Petri dish with 5 ml of E2 medium and a magnetic stir bar placed on a magnetic stirrer plate inside the incubator, as described above. Videos samples were later used for offline data recovery using ImageJ 1.48v software (National Institutes of Health, Bethesda, USA) and MTrackJ (Biomedical Imaging Group Rotterdam, Rotterdam, The Netherlands). Larvae were individually tracked and their x-y coordinates at every time point were subsequently used to calculate motion values, body orientation and position relative to the rotation axis of the magnetic stir bar, which corresponded in all cases to the center of the Petri dish. Motion values were expressed as either speed (mm per second) or distance swum every 5 or 30 seconds. To quantify the proportion of larvae engaged in rheotaxis, we measured the proportion of larvae directly facing the oncoming current 120 s after the onset of vortex flows. For this we measured – three times every 10 s – the angle formed between a larva’s body axis and a line connecting the center of its head and the rotation axis of the magnetic stir bar. A larva was considered to be engaged in rheotaxis when the coefficient of variation arising from the three angles measured over 30 s remained lower than 10 % and, at the same time, it exhibited minimum body displacements, i.e., shorter than 0.5 mm * (10 ms)^−1^. To assess the average change in orientation after a swim bout (δ body angle, in degrees), we measured, as before, the angle formed between a larva’s body axis and a line connecting the center of its head and the rotation axis of the magnetic stir bar, every 933 ms over a 5 s period 120 s after the onset of vortex flows. The resulting ‘δ body angle’ values (in degrees) were then calculated as the average difference between the consecutive angles for each larva.

### General experimental design and statistical analyses

In all experiments, replicates were randomly assigned with a treatment using a computer-generated random number sequence. Also, we used blinding. For cortisol measurements, a first experimenter executed the treatments and collected and labelled the samples. A second experimenter extracted cortisol from the labelled samples and assigned them with new labels. The first experimenter then quantified cortisol using the newly encoded samples. The results were compared by the first, second and a third experimenter at the end of the blind procedure.

For behavioural measurements, a first experimenter performed the video-recordings and labelled the resultant test videos. A second experimenter then scored the behaviour of single larvae in each labelled video. Thus all videos were assessed blind. At the end of the blind procedure, the identities of the larvae were linked to the treatments and the results examined collectively in the presence of a third experimenter. In our study, all data are shown as single measurement points or mean and standard error of the mean. No data points or samples were excluded from analysis. Our sample sizes are generally employed in the field and based on previous publications^13,45^, statistical methods to predetermine sample size were not necessary. The sample sizes in Figs. 1b, 1c (top and bottom), 1d and 2a (top and bottom) are unbalanced due to the scale of the overall study (only partially shown) and the ensuing number of simultaneous groups. In our study, several experiments were being carried out in parallel. As a result, the amount of effort that could be devoted to each one of them varied over time. We therefore set up a minimum sample size for each dataset from the very beginning, based on our previous publications and experience. Whenever likely, we added as many data points as possible, depending on the parallel experiments. Data points could have been randomly selected so as to keep sample size constant across all groups. Yet the unbalanced samples did not preclude the validity of the applied tests, they constituted independent samples and pairwise comparisons remained valid in all cases. We decided against the exclusion of data points to favour a better estimate of the coefficient of variation of the various samples. The values in Figs. 1c (top) and 1c (bottom) arise from the same type of treatment, but do not necessarily involve the same set of animals. Fig. 1c top depicts a fraction (percentage) of larvae from a group that involved several trials; here sample size refers to the total number of larvae in several dishes across trials. By contrast, sample size in Fig 1c bottom refers to individuals. Normality was tested using the Shapiro–Wilk test. Normally distributed data were then tested for homogeneity of variance using Bartlett’s tests. For two-group comparisons, we used Mann-Whitney tests (for pairwise comparisons with at least one non-normally distributed dataset) and Student’s *t*-tests (two-tailed) (normally distributed data). For multiple group comparisons we used Chi-square tests, as well as Kruskal-Wallis tests and ANOVAs, followed by Dunn’s multiple comparison tests and Bonferroni’s *post hoc* tests, respectively. Analyses were made with MS-Excel (Microsoft Corp; Redmond, WA, USA), Prism 5 (Graphpad Software Inc, San Diego, CA, USA), ImageJ (Freeware) and VirtualDub (Freeware). The figures were assembled in Illustrator (Adobe).

## Data accessibility

The data that support the findings of this study are available from the authors on request.

## Acknowledgments

We thank T. Thiemann and L. Flores-García for assistance with the experiments. This work was supported by the Max Planck Society, the University Medical Center of the Johannes Gutenberg University Mainz, the German Federal Office for Education and Research (Bundesministerium für Bildung und Forschung, grant 01GQ1404) and the German Research Foundation (DFG, grant A04-CRC 1193). We thank R. Singer and A. Schoell for expert fish care.

## Author Contributions

Conceptualization, R.J.D.M. and S.R.; Methodology, R.J.D.M. and S.R.; Investigation, L.A.C-R., S.R. and R.J.D.M.; Writing – Original Draft, L.A.C-R and R.J.D.M.; Writing – Review & Editing, R.J.D.M.

